# β-HPV 8E6 Dysregulates the Hippo Signaling Pathway and Induces Aneuploidy

**DOI:** 10.1101/760439

**Authors:** Dalton Dacus, Tristan X. McCallister, Celeste Cotton, Elizabeth Riforgiate, Nicholas A. Wallace

## Abstract

Beta genus human papillomaviruses (β-HPVs) are associated with cutaneous squamous cell carcinomas (cSCCs) in a subset of immunocompromised patients. Although β-HPVs are not necessary for tumor maintenance, they are hypothesized to destabilize the genome in the early stages of cancer development. Supporting this idea, β-HPV’s 8E6 protein attenuates p53 accumulation after failed cytokinesis. This paper identifies the mechanism of this abatement. We show β-HPV 8E6 dysregulates the Hippo signaling pathway (HP). It increases pro-proliferative gene expression, enhances TEAD activity and promotes cell growth. β-HPV 8E6 also reduces LATS activation and p53-mediated apoptosis following unsuccessful division of mitotic cells. These phenotypes are dependent on β-HPV 8E6 binding and destabilizing a cellular histone acetyltransferase, p300. Despite circumventing apoptosis, β-HPV 8E6 caused increased senescence after unsuccessful cytokinesis. We linked this lack of growth to the viral protein’s inability to prevent cytoplasmic sequestration of the HP transcription factor, YAP. We also show that increased telomerase reverse transcriptase activity (a common alteration in cSCCs) acts synergistically with β-HPV 8E6 to promote cellular proliferation after abortive cytokinesis. While β-HPV 8E6 promoted aneuploidy on its own, this genome destabilization is amplified in cells that do not divide after mitosis. Although our group and others have previously described inhibition of DNA repair, to the best of our knowledge this marks the first time that a β-HPV protein has been connected to chromosome level changes in the cellular genome. This represents a substantial escalation in the known genome destabilizing properties likely to occur during a β-HPV infection.

**IMPORTANCE:** There is mounting evidence that β-HPVs contribute to cSCCs development in immunocompromised populations. They may also augment UV’s mutagenic potential, increasing cancer risk in the general population. We demonstrate that β-HPV 8E6 dysregulates the Hippo signaling pathway (HP). HP regulates cell growth and apoptosis in response to a myriad of stimuli, including failed cytokinesis. β-HPV 8E6 attenuates phosphorylation of the HP kinase, LATS, decreasing some but not all downstream signaling events. This allows binucleated cells to avoid apoptosis, however they succumb to senescence. We show that β-HPV 8E6 synergizes with a common cSCC mutation (telomerase activation) to avoid both apoptosis and senescence. We did not find any telomerase immortalized β-HPV 8E6 expressing cells that were not aneuploid after aberrant cytokinesis. This represents a substantial escalation in β-HPV E6’s known mutagenic potential.

## INTRODUCTION

The human papillomavirus (HPV) family includes over 200 double-stranded DNA viruses that are divided into five genera, all of which infect human epithelia (1). Upon infecting mucosal or cutaneous tissue, members of each genera can cause a broad array of pathologies. Of these, the most prominent diseases are the anogenital and oropharyngeal carcinomas caused by alpha genus HPVs (2, 3). Cutaneous beta genus HPVs (β-HPVs) have also been linked to tumorigenesis via high viral DNA loads in cutaneous squamous cell carcinomas (cSCCs) of immunocompromised patients, primarily in areas of the skin exposed to the sun (4–6).

β-HPV infections are common in the general population, but their contribution to cSCCs is less clear in immune competent individuals. The main aetiological factor in skin cancer pathogenesis is UV. Further, characterizations of cSCCs in the general population do not find continued β-HPV expression (7–9). Viral loads decrease as lesions progress from precancerous actinic keratosis (AK) lesions to cSCCs (10–12). These data have led to the hypothesized “hit and run” mechanism of oncogenesis, where β-HPVs cooperate with UV to enhance genomic instability in the early stages of carcinogenesis (10, 13, 14). The elevated mutational load then increases the chances of tumor progression independent of continued viral gene expression.

While it is hard to prove the role of a transient viral infection in a persistent cancer, β-HPVs are also a common resident of our skin and frequently found in AKs. Despite the billions of dollars spent on sun care products annually, 58 million Americans still have one or more AKs. Moreover, over $1 billion is spent during 5.2 million outpatient visits each year for AK treatment (15, 16). This cost, along with the emotional toll increases if these lesions develop into malignancies. Within 1 year of diagnosis an estimated 0.6% of AKs progress to cSCCs. This progression expands to 2.6% of AKs 5 years after diagnosis (17). Thus, it is important to broadly understand how these widespread infections alter the cell’s ability to maintain genome stability.

A great deal is known about the tumorigenic potential of β-HPV proteins, particularly the E6 protein. The E6 putative oncogene from β-HPV 8 (β-HPV 8E6) is enough to cause cancers in mice without UV exposure (18, 19). β-HPV 8E6 inhibits differentiation and promotes proliferation by targeting the NOTCH and TGF-β signaling pathways (20). Another central theme of β-HPV E6 proteins binding the cellular histone acetyltransferase p300, has emerged (21–24). β-HPV 8E6 and the E6 from β-HPV 5 (β-HPV 5E6) bind p300 strongly, leading to its destabilization and decreasing DNA damage repair (DDR) gene expression (22, 25, 26). β-HPV type 38 E6 (HPV38 E6) has a lower p300 binding affinity and cannot destabilize the cellular protein (27). Nevertheless, binding p300 is essential for HPV38-induced immortalization of human foreskin keratinocytes (HFKs). This suggests that p300 binding may be a shared factor in β-HPV promoted oncogenesis (28). Because p300 is a master regulator of gene expression (29, 30), there are likely other signaling pathways altered by β-HPV 8E6’s destabilization of the histone acetyltransferase.

Approximately 10% of skin cells do not divide after entering mitosis (25, 31). β-HPV 8E6 allows these cells to remain proliferative by preventing p53 stabilization in a p300-dependent manner (25). p53 accumulation requires the activation of LATS, a kinase in the Hippo signaling pathway (HP) (32). It also prevents growth by inhibiting the pro-proliferative activity of YAP/TAZ (32–34). We hypothesize that β-HPV 8E6 dysregulates the HP after aborted cytokinesis via p300 destabilization, causing a decrease in p53 levels and an increase in YAP/TAZ driven cell growth.

Our analysis of transcriptomic data from cell lines with or without decreased p300 expression identified canonical HP and downstream HP-activated pro-proliferative genes that are negatively regulated by p300. We confirm this in primary cell culture and show that β-HPV 8E6 hinders LATS phosphorylation and prevents p53-induced apoptosis after cytokinesis failure. Senescence instead prevents the long-term expansion of cells recovering from becoming binucleated. However, telomerase reverse transcriptase (*TERT*) activation acts synergistically with β-HPV 8E6 to avoid apoptosis and senescence allowing these aberrant cells to proliferate. Consistent with enhanced tolerance of aberrant cytokinesis, β-HPV 8E6 increases aneuploidy.

## METHODS

### Cell Culture

U2OS cells were maintained in DMEM supplemented with 10% FBS and penicillin-streptomycin. Primary HFKs were derived from neonatal human foreskins. HFKs and *TERT*-immortalized HFKs (obtained from Michael Underbrink, University of Texas Medical Branch) were grown in EpiLife medium supplemented with calcium chloride (60 μM), human keratinocyte growth supplement (ThermoFisher Scientific), and penicillin-streptomycin. HPV genes were cloned, transfected, and confirmed as previously described (25).

### Proliferation Assays and H2CB Cell Viability Assays

Cells were counted and 4.0 × 10^4^ cells were plated into 6 wells per cell line of 6-well tissue cultures dishes. One well was trypsinized, resuspended and counted 3 times via hemocytometer with trypan blue. For dihydrocytochalasin B (H2CB) cell viability assays, cells were grown for 24 h then treated with 2/4 μM of H2CB, re-administering fresh H2CB every 2 days while cells were trypsinized and counted 3 times via hemocytometer with trypan blue.

### RT-qPCR

Cell were lysed using Trizol (Invitrogen) and RNA isolated with the RNeasy kit (Qiagen). Two micrograms of RNA were reverse transcribed using the iScript cDNA Synthesis Kit (Bio-Rad). Quantitative real time-PCR (RT-qPCR) was performed in triplicate with the TaqMan FAM-MGB Gene Expression Assay (Applied Biosystems) and C1000 Touch Thermal Cycler (Bio-Rad). The following probes (Thermo Scientific) were used: ACTB (Hs01060665_g1), STK4 (Hs00178979_m1), LATS2 (Referred to as LATS in the text) (Hs01059009_m1), YAP1 (Hs00902712_g1), CTGF (Hs00170014_m1), CYR61 (Hs00155479_m1).

### Immunoblotting

After being washed with ice cold PBS, cells were lysed with RIPA Lysis Buffer (VWR Life Science) supplemented with Phosphatase Inhibitor Cocktail 2 (Sigma) and Protease Inhibitor Cocktail (Bimake). The Pierce BCA Protein Assay Kit (Thermo Scientific) was used to determine protein concentration. Equal protein lysates were run on Novex 4-12% Tris-Glycine WedgeWell Mini Gels (Invitrogen) and transferred to Immobilon-P membranes (Millipore). Membranes were then probed with the following primary antibodies: p53 (Calbiochem, OP43-100UG), HA-tag (Cell Signaling Technologies 3724S), GAPDH (Santa Cruz Biotechnologies sc-47724), LATS2 (Referred to as LATS in the text, Cell Signaling Technologies D83D6), Phospho-LATS1/2 (Referred to as pLATS in the text, Ser909) (Cell Signaling Technologies #9157), YAP (Cell Signaling Technologies 4912S), Phospho-YAP (Ser127) (Cell Signaling Technologies 4911S), C23 (Nucleolin) (Santa Cruz Biotechnologies sc-8031 HRP), MST2 (Cell Signaling Technologies 3952S), 14-3-3 Theta/Tau (Millipore Sigma T5942-.1MG). After exposure to the matching HRP-conjugated secondary antibody, cells were visualized using SuperSignal West Femto Maximum Sensitivity Substrate (Thermo Scientific).

### cBioPortal and Gene Ontology Analysis

Software from (www.cbioportal.org) was used to recognize, analyze, and categorize mutations and trasnscriptomic data from over 1000 cancer cells lines (35–37) and cutaneous squamous cell carcinomas (38, 39). Gene Ontology enRIchment anaLysis and visuaLizAtion tool (GOrilla) identified and visualized enriched GO terms from these data (40, 41). Analysis of the squamous cell carcinoma samples was done at (http://geneontology.org/) powered by Protein ANalysis THrough Evolutionary Relationships (PANTHER) (42, 43). The Kyoto Encyclopedia of Genes and Genomes (KEGG) was used to identify genes specific to the Hippo signaling pathway (hsa04390).

### Senescence-associated β-galactosidase Staining

Cells were seeded onto 3 6-well plates and were grown for 24 h. Then, they were treated with 4 μM of H2CB for stated times then cells were fixed and stained for senescence-associated β-galactosidase (β-Gal) expression according to the manufacturer’s protocol (Cell Signaling Technologies).

### Immunofluorescence Microscopy

Cells were seeded onto either 96-well glass bottom plates (Cellvis) or coverslips and grown overnight. Cells treated with H2CB for specified time and amount of H2CB were fixed with 4% formaldehyde. Then 0.1% Triton-X solution in PBS was used to permeabilize the cells, followed by blocking with 3% bovine serum albumin in PBS for 30 minutes. Cell were then incubated with the following: p53 (Cell Signaling Technologies 1C12), YAP (Cell Signaling Technologies 4912S), Ki67 (Abcam ab15580), alpha tubulin (Abcam ab18251). The cells were then washed and stained with the appropriate secondary antibodies: Alexa Fluor 594 goat anti-rabbit (Thermo Scientific A11012), Alexa Fluor 488 goat anti-mouse (Thermo Scientific A11001). Washed with PBS 2X and stained with 28 μM DAPI in PBS for 12 min before a final wash in PBS and visualization with Zeiss LSM 770 microscope. Foci were analyzed using ImageJ previously described in (44).

### Luciferase Reporter Assays

Reporter assays were performed using Dual-Luciferase Reporter Assays System (Promega). Transfected cell lysates with 8xGTIIC-luciferase, a gift from Stefano Piccolo (Addgene plasmid #34615; (http://n2t.net/addgene:34615); RRID:Addgene_34615) were prepared in 100 μl of passive lysis buffer 48 h post transfection and 40 μl of lysate was used for each reading. Readings were done in triplicate using GloMax Navigator Microplate Luminometer (Promega) and values were normalized for transfection efficiency using co-transfected renilla luciferase plasmid.

### Apoptosis Assay

After H2CB treatment, HFKs were harvested via trypsinization then counted while incubating at 37°C for 30 min. After incubation cells were resuspended to 1 × 10^6^ cells/ml. Next, cells were stained with 100 μg/ml of propidium iodide and 1X annexin-binding buffer following the protocol from Dead Cell Apoptosis Kit (Invitrogen V13242). Stained cells were imaged with the Countess II FL Automated Cell Counter (Invitrogen). Images were processed using ImageJ software.

### Sub-Cellular Fractionation

Cells were seeded at 5.0 × 10^5^ cells/10 cm^2^ plate and grown for 24 h. Cells were then treated with 4 μM H2CB for 3 days, then washed with PBS and recovered in fresh EpiLife for 3 days (After H2CB treatment). Before and after H2CB exposure, cells were then washed with ice-cold PBS and divided into cytosolic and nuclear fractions via Abcam’s subcellular fractionation protocol. Afterwards lysates were treated the same as in the Immunoblot section.

### H2CB Recovery Assay

Cells were counted and 1.0 × 10^5^ cells were seeded on a 10 cm^2^ tissue culture plate and grown for 24 h. Cells were then treated with 4 μM H2CB, refreshing the H2CB every 3 days. After 3 or 6 days, the cells were washed with PBS and given fresh EpiLife. Once cells reached 90% confluency they were counted and 9.0 × 10^4^ cells were reseeded. This process was continued for 30 days if possible.

### Chromosome Counts via Metaphase Spread

Before and after H2CB exposure, *TERT*-immortalized HFKs were grown to 80% confluency then chromosomes were detected and counted as previously described (45).

### Statistical Analysis

Unless otherwise noted, statistical significance was determined by a paired Student *t* test and was confirmed when appropriate by two-way analysis of variance (ANOVA) with Turkey’s correction. Only *P* values less than 0.05 were reported as significant.

## RESULTS

### β-HPV 8E6 increases TEAD responsive genes in a p300-dependent manner

Animal models show that certain β-HPV E6 genes can contribute to UV’s carcinogenesis (18, 19, 23). *In vitro* studies from our group and others have added molecular details by describing β-HPV E6’s ability to impair the DDR by destabilizing p300 (18, 22, 27, 28, 46). However, β-HPV E6’s oncogenic potential probably extends beyond DDR inhibition, as it can induce skin tumors in mice without UV exposure (18). Further, p300 is a master transcriptional regulator, meaning the impact of its destabilization is unlikely to be confined to DDR pathways (29, 30). There are DDR-independent pathways that protect genome fidelity, whose inhibition would be consistent with the “hit and run” model by which β-HPV infections are believed to contribute to cSCCs (47–49). To identify p300-regulated pathways that could contribute to β-HPV E6-associated genome destabilization, we performed an *in silico* screen by comparing RNAseq data among 1020 cancer cell lines grouped by their relative p300 expression levels, mimicking β-HPV 8E6 expression (Supplementary Data 1) (35, 50, 51). We identified genes that were upregulated (p-value<0.05) when p300 was decreased (z-score<−1.64). Next, gene ontology (GO) analysis was performed using GOrilla to identify pathways that were significantly altered when p300 expression was reduced (52, 53). This revealed significant up-regulation of genes involved in the HP (Fig. 1A), specifically TEAD1/4 which are transcriptional activators that promotes cell proliferation and restrict apoptosis (54, 55). We then performed a more detailed analysis of the 154 HP genes using KEGG to define the pathway. When p300 expression was reduced, many canonical HP genes were up-regulated. However, the most striking changes occurred in pro-proliferative TEAD-responsive genes (CYR61, CTGF, AXL, SERPINE1) (Fig. 1B).

**Figure 1:**
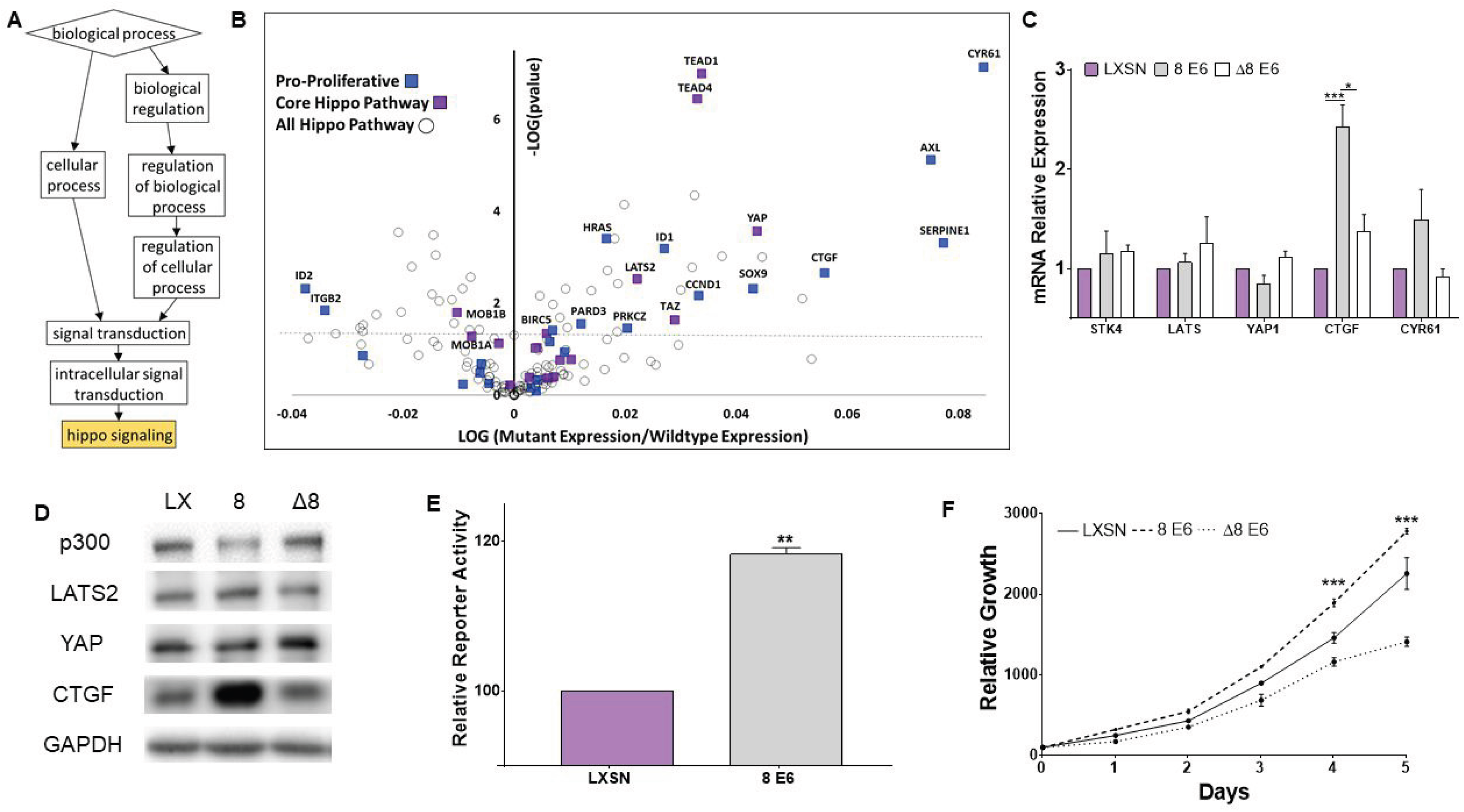
β-HPV 8E6 alters Hippo pathway signaling in a p300-dependent manner. (A) Gene ontology of 1020 cancer cell lines via GOrilla. Boxes show GO biological processes terms. Boxes descend from general to specific functions. Gold color indicates p ≤ 0.001. (B) Volcano plot of 154 HP genes in 1020 cancer cell lines with decreased EP300 expression. The colors blue and purple label pro-proliferative TEAD targets and core HP genes, respectively. The horizontal line denotes p = .05. (C) HP and TEAD-regulated genes mRNA expression in HFKs measured by RT-qPCR and normalized to β-actin mRNA. (D) Representative immunoblots of HP proteins in HFKs LXSN, β-HPV 8E6 and β-HPV Δ8E6. (E) TEAD-responsive promoter activity in *TERT*-HFKs. (F) Relative growth recorded over 5-day period. Figures depict means ± standard error of mean. n ≥ 3. * denotes p ≤ 0.05, ** denotes p ≤ 0.01, *** denotes p ≤ 0.001.

Because β-HPV infect the cutaneous epithelia, we chose HFKs as a relevant *in vitro* model for validating our computational data. Vector control (LXSN), β-HPV 8E6 expressing HFKs were generated. To confirm p300 dependence, HFKs expressing β-HPV Δ8 E6 (residues 132–136 were deleted) were also generated. This mutant can no longer bind or destabilize p300 (56). RT-qPCR indicated that β-HPV 8E6 did not significantly change the expression of HP genes (Fig. 1C). However, two pro-proliferative TEAD-responsive genes were upregulated, CTGF and CYR61 (Fig. 1C). Although, only CTGF’s increase was statistically significant these results parallel our *in silico* analysis (Fig. 1C). To test whether this change in transcription would manifest at the protein level, we observed HP protein abundance by immunoblot. β-HPV 8E6 increased LATS and CTGF protein levels (Fig. 1D). Confirming HPV E6 expression, p300 was significantly decreased (Fig. 1D). These data suggested that β-HPV E6 increases TEAD activity, so we used a TEAD-responsive luciferase promoter (57). β-HPV 8E6 caused a slight, but significant increase in luciferase, consistent with an ability to elevate TEAD promoter activity (Fig. 1E). As a functional readout of HP activation, we defined cell proliferation rates. Compared to vector control HFKs, β-HPV 8E6 increased growth in a p300-dependent manner, over a 5-day period (Fig. 1F).

### β-HPV 8E6 increases the tolerance of failed cytokinesis in a p300-dependent manner

H2CB blocks cytokinesis via inhibiting actin polymerization, inducing tetraploidy. This activates the HP through LATS phosphorylation (32). Since we found decreased p300 expression correlated with enhanced TEAD activity, we hypothesized that β-HPV 8E6 hindered the HP’s upstream protein’s response to unsuccessful division after mitosis by binding and destabilizing p300 (Fig. 1). Our first step towards testing this hypothesis was to characterize H2CB activity in human osteosarcoma-derived U2OS and HFK cells expressing vector control, β-HPV 8E6, or β-HPV Δ8 E6. Binucleation was visualized by brightfield microscopy (Fig. 2A and 2B). H2CB universally increased the frequency of cells with more than one nucleus. Overall, there was an 8.5-fold increase in U2OS cells (8.6% ± 6.7% versus 72.7% ± 11.7%, n=3, *P* < 0.05) and a 7.45-fold increase in HFK cell lines (7.4% ± 1.4% versus 55.3% ± 5.0%, n=3, *P* < 0.05) (Fig. 2C and 2D). There was no statistical difference in H2CB-induced failed cytokinesis among the cell lines. Next, we began defining the cellular response to endoreplication without proper separation. U2OS cells were seeded at equal density and tracked for five days after H2CB treatment. As expected, inhibiting cell division was toxic with only fractions of the starting population of vector control cells remaining at the end of our time course (46.1% ± 5.6% of the cells initially seeded). β-HPV 8E6 significantly reduced H2CB’s deleterious effects (79.7% ± 12.8% of the original population) (Fig. 2E). β-HPV Δ8E6 behaved indistinguishably from wildtype cells consistent with a p300-dependent mechanism of action (56.9% ± 7.6% of the starting population) (Fig. 2E). A similar trend was seen in HFKs expressing vector control, β-HPV 8, or −Δ8E6, where β-HPV 8E6 expressing cells had reduced cell death when treated with H2CB over the course of 5 days (Fig. 2F). Using Ki-67 as a proliferation marker (58), we found that despite allowing more cells to survive H2CB, β-HPV 8E6 did not promote growth (Fig. 2G).

**Figure 2:**
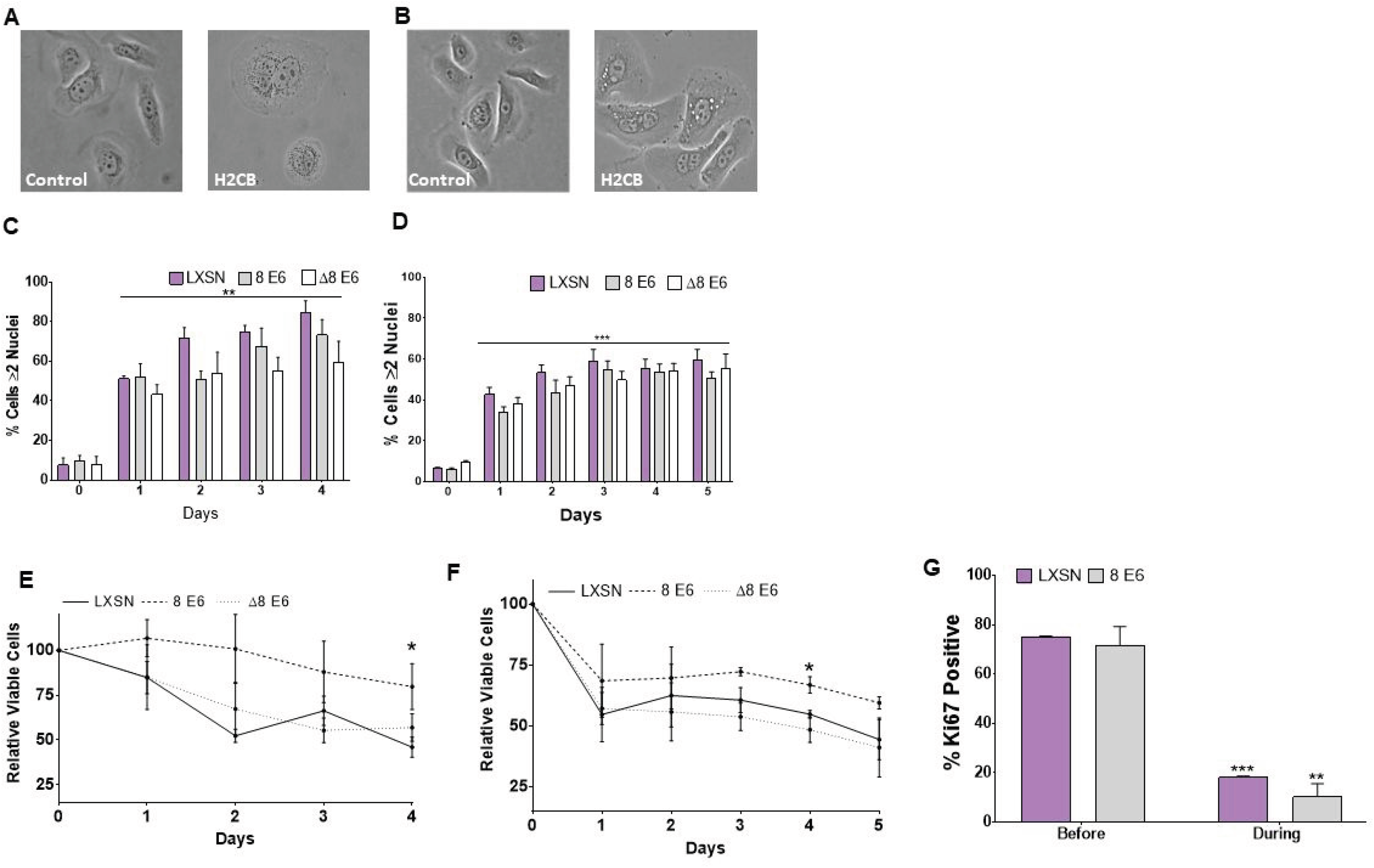
β-HPV 8E6 increases tolerance of failed cytokinesis by destabilizing p300. Representative images of (A) U2OS and (B) HFK cells with and without H2CB. Quantification of (C) U2OS and (D) HFK cells with 2 or more nuclei as a function of time in H2CB. Relative counts of (E) U2OS and (F) HFK cells treated with H2CB. Day 0 set to 100. (G) Percent of HFKs staining positive for Ki67. At least 100 cells/line were imaged in three independent experiments. Figures depict means ± standard error of mean. n ≥ 3. * denotes p ≤ 0.05, ** denotes p ≤ 0.01, *** denotes p ≤ 0.001.

### β-HPV 8E6 attenuates p53 accumulation induced by failed cytokinesis by inhibiting pLATS activation

We previously reported that β-HPV 8E6 attenuated p53 stabilization after spontaneous binucleation (25). We used immunofluorescence microscopy and immunoblotting to confirm that this phenotype was maintained in cells exposed to H2CB. β-HPV 8E6 prevented the increase in p53-staining seen in vector control cells grown with H2CB (Fig. 3A and 3B). Consistent with past observations, the frequency of p53 staining and binucleation were equivalent in vector control (59.5% ± 4.3% vs. 51.35% ± 1.4) (Fig. 2C and 3B). Validating the immunofluorescence data, immunoblot analysis confirmed that β-HPV 8E6 expressing cells limited p53 accrual when binucleated (Fig. 3C). Supporting a p300-dependent mechanism, β-HPV Δ8E6 did not suppress p53 induction (Fig. 3A-D). Interestingly, we did not see the increase in PI-positive cells after H2CB treatment that we had anticipated (Fig. 3E).

**Figure 3:**
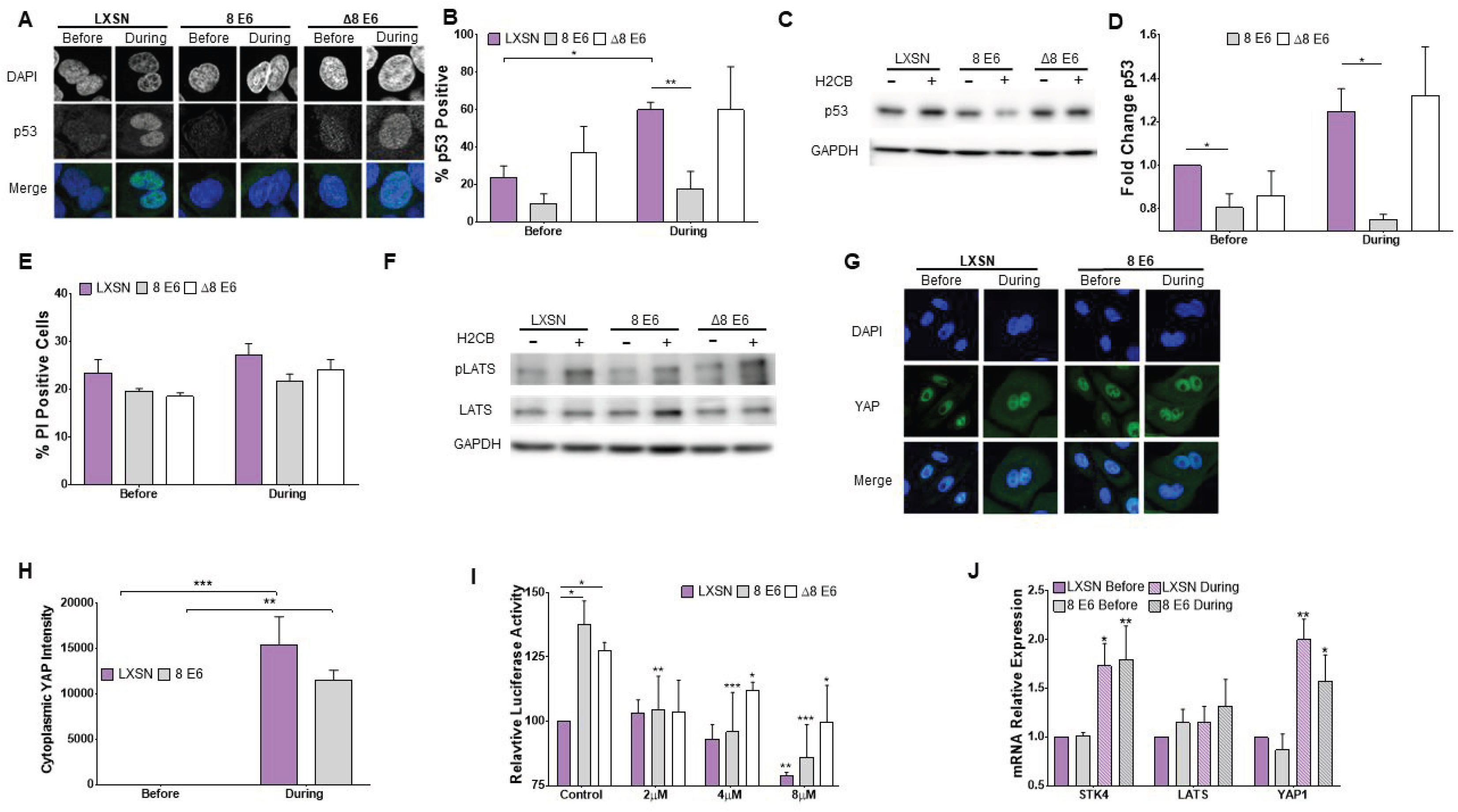
β-HPV 8E6 hinders LATS activation reducing p53 accumulation after failed cytokinesis. (A) Representative images of p53 and DAPI staining in cells with and without H2CB. (B) Percentage of p53 positive U2OS cells. (C) Representative immunoblot of p53 with and without H2CB. (D) Densitometry of immunoblots described in (C). GAPDH was used as a loading control. Data was normalized to p53 levels in untreated LXSN cells (set to 1). (E) Percentage of propidium iodide stained HFK cells with and without H2CB. (F) Representative immunoblot of pLATS and totals LATS protein levels in HFK cells with and without H2CB exposure. (G) Representative images of YAP and DAPI stained HFK cells with and without H2CB. (H) Relative cytoplasmic YAP intensity in HFK cells. Untreated cytoplasmic YAP intensity was set to 0. At least 50 cells/line were imaged from three independent experiments (I) TEAD-responsive promoter activity in U2OS cells treated with H2CB. (J) STK4, LATS, YAP1 expression before and after H2CB exposure measured by RT-qPCR n=2. Figures depict means ± standard error of mean. n ≥ 3. * denotes p ≤ 0.05, ** denotes p ≤ 0.01, *** denotes p ≤ 0.001.

LATS activation is necessary for p53 to accumulate in tetraploid cells (32). To determine if β-HPV 8E6 reduced p53 accumulation by attenuating LATS phosphorylation, we induced failed cytokinesis and detected pLATS. Vector control cells showed a sharp increase in phosphorylated LATS (pLATS) relative to total LATS (Fig. 3F). Only total LATS levels increased when β-HPV 8E6 was expressed (Fig. 3D). The induction of LATS phosphorylation in β-HPV Δ8E6 HFKs resembled the response in control cells, consistent with a p300-dependent mechanism (Fig. 3F).

A branch point in the HP occurs with LATS activation, which stabilizes p53 and phosphorylates YAP. This sequesters YAP in the cytoplasm preventing it from promoting transcription of pro-proliferative TEAD-responsive genes (57, 59, 60). To understand the breadth of β-HPV 8E6’s HP dysregulation, we examined YAP sequestration by immunofluorescence microscopy. YAP translocated from the nucleus to cytoplasm in binucleated vector control HFKs (Fig. 3G-H). Neither β-HPV 8E6 nor β-HPV Δ8E6 changed this. Since YAP is a TEAD co-transcriptional activator, an increase in cytoplasmic YAP should result in a drop in TEAD activity (61). Utilizing the TEAD-reporter assay from Fig. 1E, we found that TEAD-driven transcription was reduced by H2CB-induced binucleation equally among the cell lines (Fig. 3I). While H2CB treatment significantly increased expression of STK4/MST1 and YAP in vector control HFKs, LATS expression remained constant. β-HPV 8E6 did not alter these changes (Fig. 3J). Consistent with specific inhibition of LATS activation, β-HPV 8E6 did not change the abundance of the other HP proteins (Supplemental Fig. 1A).

### β-HPV 8E6 limits apoptosis in cells recovering from failed cytokinesis

Elevated p53 levels induce apoptosis in cells that do not divide after mitosis (32, 62). β-HPV 8E6’s ability to reduce H2CB toxicity and diminish p53 accumulation, suggests that it limits apoptosis (Fig. 2E-F and 3A-D). However, we did not see increased apoptosis in any of the cells grown in H2CB for 24 hours (Fig. 3D). We hypothesized that apoptosis plays a larger role in rebounding from aberrant cytokinesis. To test this, we observed cells after H2CB was removed. There were no significant changes in binucleation frequency in vector control HFKs. However, β-HPV 8E6 caused a significant decrease in supernumerary nuclei frequency (59.5% ± 1.0% vs 36.3% ± 1.3%) (Fig. 4A). β-HPV Δ8E6 HFKs remain binucleated like vector control cells. (Fig. 4A). Qualitative observations indicated that LXSN HFKs were dying and or becoming senescent. In contrast, β-HPV 8E6 expression allowed cells to undergo another round of replication, after which they frequently became mononucleated and heavily vacuolated (Supplementary Fig. 1B).

**Figure 4:**
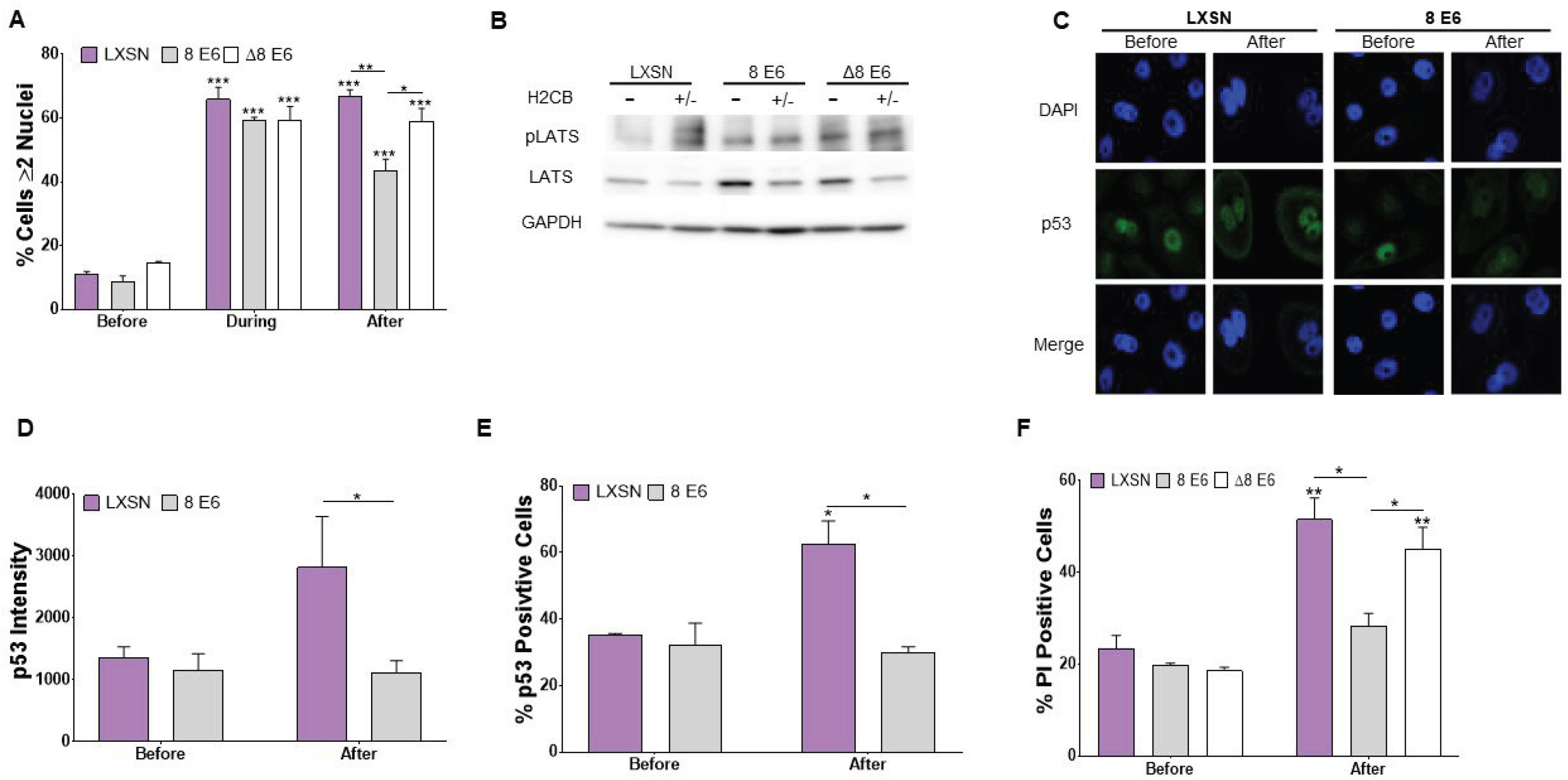
β-HPV 8E6 lowers the frequency of apoptosis in cells with two nuclei. (A) Percent of HFK cells with ≥ 2 nuclei per cell before, during and after H2CB. (B) Representative immunoblots of pLATS and totals LATS in HFKs before H2CB and after H2CB was removed. (C) Representative images of p53 and DAPI staining in cells before and after H2CB. (D) Mean p53 intensity in HFK cells before and after H2CB. At least 200 cells/line were imaged across three independent experiments. (E) Percent of HFK cells that stained positive for p53 before and after H2CB. (F) Percentage of propidium iodide stained HFK cells before and after H2CB. Figures depict means ± standard error of mean. n ≥ 3. * denotes p ≤ 0.05, ** denotes p ≤ 0.01, *** denotes p ≤ 0.001.

To determine if β-HPV 8E6 retained its ability to suppress LATS, we observed cells when H2CB was removed. Immunoblots demonstrated that β-HPV 8E6 continued to attenuate the normal LATS phosphorylation (Fig. 4B). Similarly, immunofluorescence microscopy found β-HPV 8E6 decreased both the frequency and intensity of p53 staining (Fig. 4C-E). There was a notable similarity between the frequency of p53 staining and binucleation as cells recovered (Fig. 4A, D-E). As we predicted, apoptosis was a typical response to H2CB removal that paralleled both p53 induction and binucleation (Fig. 4A and E-F). β-HPV 8E6, but not β-HPV Δ8E6, decreased apoptosis after H2CB (Fig. 4F). Thus, β-HPV 8E6’s attenuation of p53 accumulation minimized cell death in response to abortive cytokinesis.

### β-HPV 8E6 increases senescence during resolution of failed cytokinesis

Despite avoiding apoptosis, β-HPV 8E6 did not completely absolve cells from the consequence of failed cytokinesis. Cell morphology, stifled proliferation, and reduced apoptosis led us to suspect that β-HPV 8E6 increased senescence (63, 64). We used Ki67 staining to define proliferation. β-HPV 8E6 had markedly lower Ki67 intensity during and after H2CB treatment (Fig. 5A and 5B 572.0 ± 183.6 and 699.1 ± 278.6 respectively). We used senescence-associated β-Galactosidase (SA β-Gal) activity as a marker of senescent cells (65). β-HPV 8E6 significantly increased the frequency of SA β-Gal staining after H2CB was removed (52.0% ± 7.0% vs 78.5% ± 6.8%) (Fig. 5C-D). Immunofluorescence microscopy and subcellular fractionation show that post-H2CB, cytoplasmic YAP sequestration was analogous among our cell lines, meaning this branch of the HP remained in tacked, despite muted the LATS phosphorylation caused by β-HPV 8E6 (Fig. 5E-5G). These data are consistent with β-HPV 8E6 allowing cells to escape apoptosis only to “get caught” by senescence.

**Figure 5:**
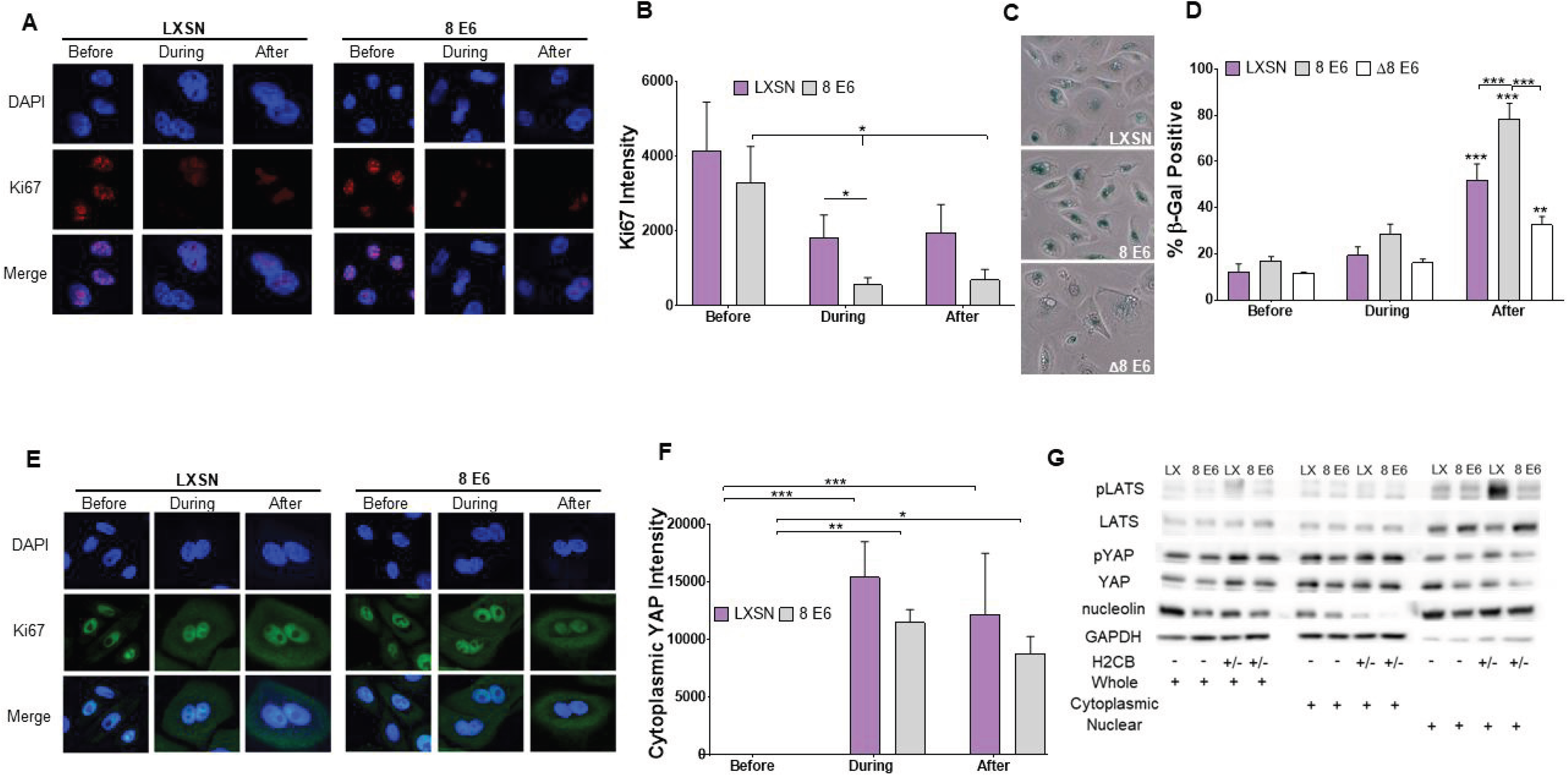
β-HPV 8E6 increases senescence in cells recovering from failed cytokinesis. (A) Representative immunofluorescence microscopy images of HFK cells before, during and after H2CB. Ki67 staining marks proliferating cells. DAPI marks nuclei. (B) Average Ki67 intensity of ≥150 images of HFK cells before, during and after H2CB. (C) Representative images of HFK cells stained for β-Gal activity (blue). (D) Quantification of β-Gal positive HFK LXSN, β-HPV 8E6 and Δ8E6 cells before, during and after H2CB exposure. (E) Representative images of HFK cells stained for YAP and with DAPI. (F) Cytoplasmic YAP intensity in HFK cells before, during, and after H2CB treatment (≥ 205 images from 3 independent experiments). (G) Subcellular fractionation of HFKs harvested before and after H2CB. Hippo pathway proteins were probed via immunoblot. GAPDH and nucleolin serve as cytoplasmic and nuclear loading controls, respectively. Figures depict means ± standard error of mean. n ≥ 3. * denotes p ≤ 0.05, ** denotes p ≤ 0.01, *** denotes p ≤ 0.001.

### *TERT* allows cells to bypass senescence and recover after H2CB-induced failed cytokinesis

Curious as to whether mutations common in cSCC were associated with bypassing senescence, we queried sequencing data from 68 cutaneous cSCC samples (38, 39). First, we identified mutated genes in cSCCs and ranked them by their frequency. The top 10% of these genes were analyzed using the web-based gene ontology software, PANTHER (Supplementary Data 2) (42, 43, 66) Proliferation and senescence were identified as biological processes commonly altered in cSCCs and expected to limit senescence. Of the relevant genetic changes, *TERT* activation was selected because it was independently verified to be common in cSCCs (67–70). We used previously characterized keratinocytes immortalized by telomerase activation and transduced with HA-tagged β-HPV 8 E6 or vector control (TERT-HFK LXSN and TERT-HFK β-HPV 8E6) (71, 72).

To determine if β-HPV 8E6 and *TERT* activation synergize, we exposed cells to H2CB for 3 days and monitored their recovery over time after the drug was removed. Although β-HPV 8E6 could mitigate some of H2CB’s short term toxicity, we were never able to get HFKs to grow for more than a few passages after H2CB. In contrast, *TERT* activation alone and in combination with β-HPV 8E6 allowed them to sustain growth after 3-days of H2CB treatment (Fig. 6B). Binucleation was increased by *TERT* activation with and without β-HPV E6 (51.4% ± 3.5% vs 87.9% ± 2.9%), (Fig. 6C). SA β-Gal staining was increased in all *TERT*-HFKs exposed to H2CB (19.6% ± 3.6% vs 63.4% ± 3.8%),(Fig. 6D). Of the senescent cells, the fraction that had ≥2 nuclei increased in *TERT*-HFKs (19.2% ± 0.9% vs 88.8% ± 3.7%), (Fig. 6E).

**Figure 6:**
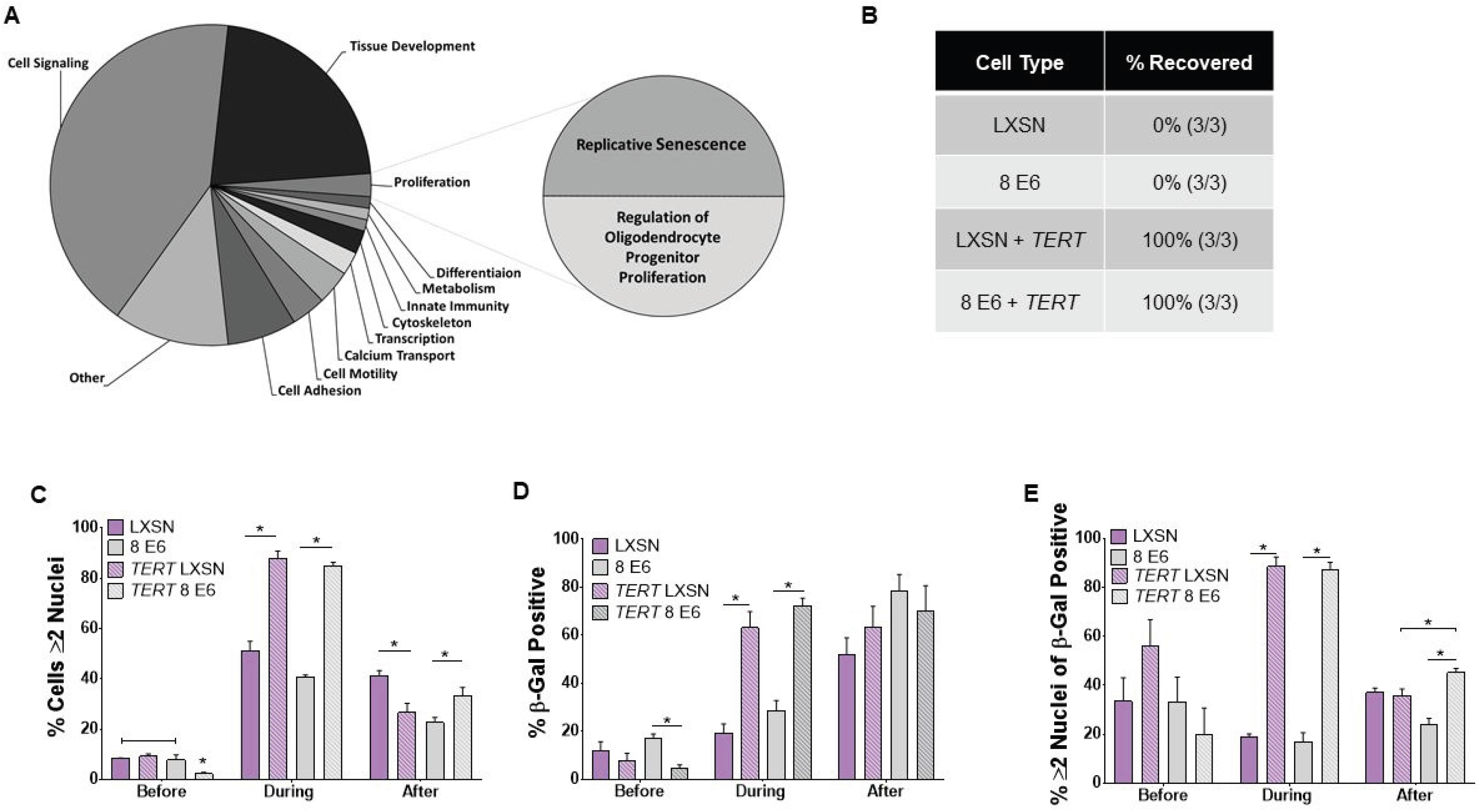
*TERT* expression promotes senescence bypass and recovery after failed cytokinesis. (A) GO analysis of common mutations in cSCCs analyzed with PANTHER software. (B) Percentage of HFKs capable of long-term growth after 3 days H2CB exposure. (C) Quantification of cells with more than 1 nucleus before, during, and after H2CB exposure. (D) Quantification of β-Gal staining in HFK cells before, during and after H2CB. (E) Percentage of cells with ≥2 nuclei that were β-Gal positive before, during and after H2CB. Figures depict means ± standard error of mean. n ≥ 3. * denotes p ≤ 0.05, ** denotes p ≤ 0.01, *** denotes p ≤ 0.001.

### β-HPV 8E6 and *TERT* activation synergize, allowing genetically unstable cells to proliferate

Although telomerase activation alone allowed cells to recover from failed cytokinesis, we wanted to determine if this ability could be augmented by β-HPV 8E6 after extended H2CB treatment. To this end, we grew cells in H2CB for an additional 3 days. The longer exposure resulted in a notably lower *TERT*-HFK LXSN cell survival (Fig. 7A). In contrast, *TERT*-HFK β-HPV 8E6 cells rebounded every time (Fig. 7A). When both cell lines lived, β-HPV 8E6 expression made them resume normal growth rates sooner (Fig. 7B). We hypothesized that surviving had high costs regarding genome stability and tested this idea by examining changes in ploidy (Fig. 7C). After H2CB exposure, polyploidy and aneuploidy dominated, with most cells having ~96 chromosomes (Fig. 7D-E). Chromosome abnormalities were further exacerbated by β-HPV 8E6 with and without H2CB (Fig. 7D-E).

**Figure 7:**
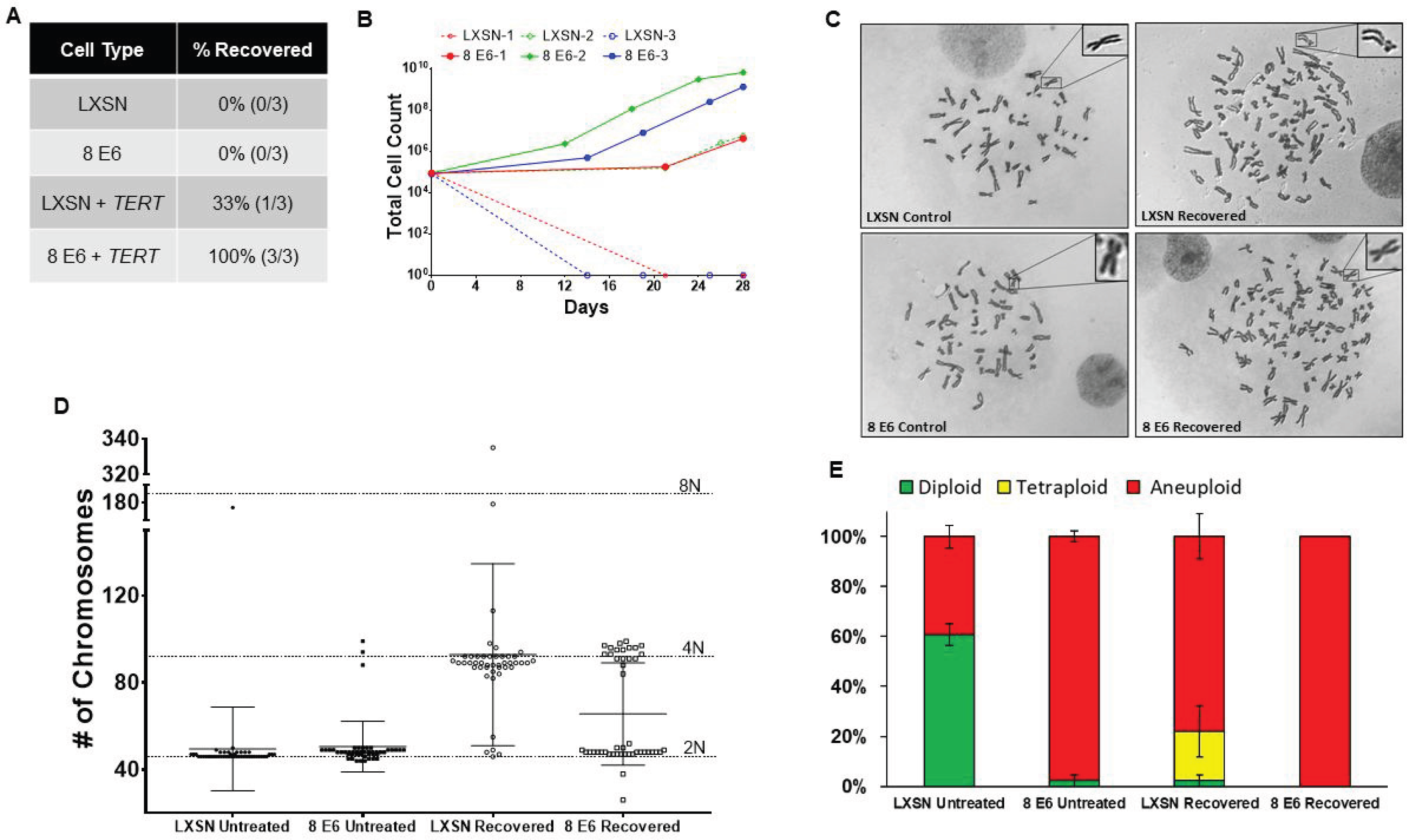
β-HPV 8E6 and *TERT* expression synergistically promote genome instability. (A) Percentage of HFK cells capable of long-term growth after 6 days in H2CB. (B) Growth curves for HFK cells after 6 days of H2CB. (C) Representative images of metaphase spreads. Insert on top right corner shows magnification. (D) Chromosome counts from at least 45 cells before or after 6 days of H2CB. (E) Relative frequency of diploidy (green), tetraploidy (yellow), and aneuploidy (red) before and after 6 days of H2CB exposure. Figures depict means ± standard error of mean. n ≥ 3.

## DISCUSSION

The proliferation of tetraploid cells stemming from mitotic failures threatens genomic stability and enable tumor development by causing aneuploidy (47, 73). The HP minimizes this risk by halting growth and initiating apoptosis (32, 74). Mitotic skin cells face a 10% risk of becoming tetraploid, thus, the HP is particularly important for maintaining their genomic stability (25, 31). β-HPV 8E6 disrupts multiple cell signaling pathways necessary for DNA repair and regulating differentiation. Much of its ability to disrupt DNA repair has been linked to its destabilization of p300. We show that the reduction of p300 also hinders the HP’s activation. Adding to the mechanisms known to promote cell growth, reduced HP activity is accompanied by slightly increased proliferation and elevated pro-proliferative TEAD activity. However, β-HPV 8E6 also allows binucleated cells to avoid apoptosis. More specifically, β-HPV E6 attenuates LATS phosphorylation leading to a blunted p53 response. While this temporarily promotes proliferation, senescence acts as a failsafe mechanism preventing extensive growth of these damaged cells. This checkpoint can be bypassed if β-HPV 8E6 is expressed in cells immortalized by telomerase activation. Consistent with dysregulation of the cell’s response to failed cytokinesis, β-HPV 8E6 increased chromosomal instability (i.e., aneuploidy).

The strongest criticism of the idea that β-HPV infections contribute to cancer, is the fact that most people get infected but significantly fewer people get cSCCs. We show that β-HPV 8E6’s ability to cause genome instability is enhanced by telomerase activation, suggesting that β-HPV infections may be more tumorigenic in certain groups of people. The same principle would apply within an individual as well. If such an “at risk” group(s) exists, mutations in the host cell’s genome at the time of infection would matter as much as whether a viral infection occurred. This could explain the current struggles of epidemiologists to link β-HPVs to skin cancers, as existing observations have largely focused on the presence/absence of β-HPV. Relevant to the work presented here, telomerase activating mutations are common during cSCC development (68, 69).

We also extend the long history of using viral oncogenes to learn about cell biology. Our data links p300 to the HP and TEAD responsive gene expression. We also confirm that LATS is activated in binucleated cells. Moreover, we show that senescence is triggered in binucleated cells that bypass apoptosis and that telomerase activity is a limiting factor in the propagation of binucleated cells. Our observation that cells sense and respond to the disruption of actin polymerization before it culminates in a binucleated cell is novel. Finally, we show that the deleterious effects of failed cytokinesis increase when resolution of the ensuing polyploidy is delayed.

Others have suggested that β-HPV influences the HP, most notably by binding PTPN14 (75). Genus α HPV oncogenes also dysregulate the pathway (76, 77). Our data adds another link between HPVs and the HP. The shared manipulation of the HP suggests HPVs gain an evolutionary advantage by abrogating the signaling cascade. The “motivation” could stem from the modest growth advantage that we report; however, this weak phenotype seems unlikely to drive convergent evolution toward HP dysregulation. Given the HP’s role in immunity, it is more enticing to speculate that targeting the HP helps HPVs avoid an immune response (78, 79). Indeed, an MST1 deficiency leads to more prevalent β-HPV infections (80). Since the HP was only discovered 14 years ago (81), there could be other currently unknown advantageous to be gained by disrupting the pathway.

## Figure Legends

**Supplemental Figure 1:**
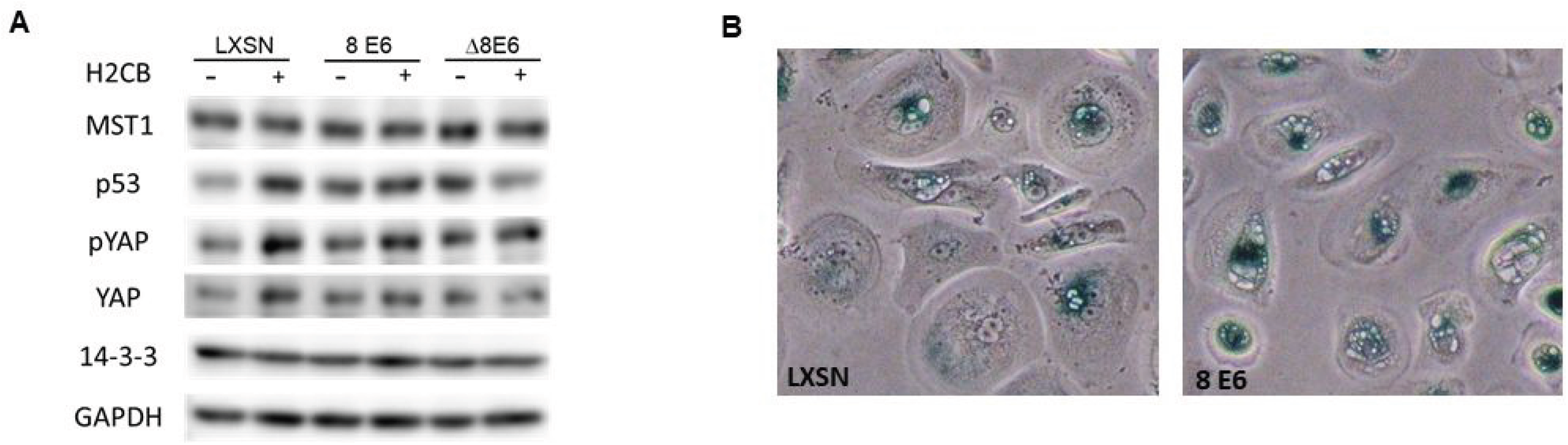
(A) Representative immunoblot core HP protein levels in HFK cells with and without H2CB exposure. (B) Representative images of HFK cells stained for β-Gal activity (blue) and visualization of cytoplasmic vacuoles.

## Acknowledgements

We thank and acknowledge the KSU-CVM Confocal Core, especially Joel Sanneman for assisting with our immunofluorescence imaging. Michael Underbrink for providing the *hTERT*-immortalized HFKs. Stefano Piccolo for gifting the 8xGTIIC plasmid. Also, we thank members of the Zhilong Yang lab for their support.

This work was supported by Department of Defense CMDRP PRCRP CA160224 (NW) and made possible through generous support from the Les Clow family and the Johnson Cancer Research Center at Kansas State University.

